# A Simple Nuclear Contrast Staining Method for MicroCT-Based 3D Histology Using Lead(II) Acetate

**DOI:** 10.1101/2020.09.18.303024

**Authors:** Brian Metscher

**Affiliations:** Department of Evolutionary Biology, Theoretical Biology Unit University of Vienna, Austria

**Keywords:** X-ray microtomography, 3D histology, staining, lead(II) acetate, hematoxylin, limb development, morphology, imaging

## Abstract

X-ray microtomography (microCT) enables histological-scale 3D imaging of many types of biological samples, but it has yet to rival traditional histology for differentiation of tissue types and cell components. This report presents prima facie results indicating that a simple lead(II) acetate staining solution can impart preferential X-ray contrast to cell nuclei. While not strictly selective for nuclei, the staining reflects local cell-density differences. It can be applied in a single overnight treatment and does not require hematoxylin staining or drying of the sample. The stain is removable with EDTA, and it may enhance early calcifications. A basic protocol is given as a guide for further testing and optimisation.

## INTRODUCTION

General contrast stains have enabled microCT imaging to become a standard tool for visualizing 3D micromorphology of soft tissues, but after a full decade, the lack of tissue specificity still limits the use of X-ray microscopy for 3D histological investigation of tissues and organisms. The need for 3D histology is widely recognised in pathology, tissue reconstruction, and other applications (Pichat et al., 2018, Roberts et al., 2012). In a recent advance toward 3D histological contrasting for microCT, Busse et al. (2018) demonstrated an X-ray contrasting method using eosin and critical point drying.

Although an enormous number of histological staining techniques have been used in the past century (Gray, 1954), the prevalent standard is hematoxylin and eosin (H&E) (Bancroft and Gamble, 2008, Wittekind, 2003). This generally applicable method most often comprises nuclear staining with hematoxylin (actually a metal-dye complex with its oxidised form, hematein) followed by a less specific counterstain, typically eosin or a similar anionic dye (Kiernan, 2008a), which together render good overall differential colouring of cellular components. Some hematoxylin solutions are prepared with a metal mordant (commonly aluminium) to form a “lake” which is applied as a stain, but in some procedures the metal is applied before the hematoxylin as a pre-mordant. In this case the metal binds to the target tissue and the hematoxylin binds to the metal to render a coloured complex (Kiernan, 2015).

Heavy metal treatments of various kinds are a mainstay in electron microscopy for imparting contrast to tissue elements (Hayat, 1970, Björkman and Hellstrom, 1965). Metal-mordanted hematoxylin methods are natural candidates for X-ray dense nuclear staining, and many such methods have been explored in the history of histology (Smith, 2010). Among these, several lead hematoxylin methods are known (Solcia et al., 1969, Guida and Cheng, 1980), but they are not frequently used in light microscopy, as they offer no particular advantages over the well-established aluminium and iron methods.

However, Müller et al. (2018) have recently demonstrated nucleus-specific contrasting in microCT imaging of mouse liver tissue using a lead-mordanted hematoxylin stain. Their results showed definite preferential staining of nuclei in liver tissue, but the procedure took about a week to complete and included critical-point drying of the sample.

Preliminary results presented here demonstrate a fast and simple method of cell-nucleus contrasting for microCT, based on the lead(II) acetate mordant used in previous methods. The present method, dubbed “Bleikern,” does not require hematein staining of the bound mordant metal and can be applied to fixed and stored tissue samples in one overnight treatment.

## METHODS

The embryo and tissue samples used were left over from previous studies, and some were of uncertain provenance. All had been fixed in paraformaldehyde (PFA) or buffered formalin (NBF). The mouse intestines were from adult wild-type BL6 mice that were NBF-fixed (as part of a study at the Medical University of Vienna), stored at 4°C in water, and dissected and re-processed as needed. Chick embryo limbs (approx. stage HH30) had been PFA-fixed and stored in methanol at −20°C. Axolotl limbs were taken from NBF-fixed specimens (ca. 20mm length) of unknown source, stored in 70% ethanol for at least 15 years. Mouse embryos and embryonic limbs were fixed and stored in NBF at 4°C for several years; these also were left over from an earlier project and their source is not known.

The staining solution was lead(II) acetate trihydrate (Pb(CH_3_COO)_2_·3H_2_O, CAS 6080-56-4; Sigma-Aldrich 467863) dissolved in distilled water at 20% w/v (0.53M, pH~5.5). This solution was stored at room temperature and could be used for at least two weeks (up to four weeks according to Müller et al. (2018)). Concentrations of 10%, 2%, and 1% were also prepared in distilled water. All samples were washed in distilled water before lead staining.

Mouse intestine samples were either stored in 70% ethanol at room temperature or were transferred directly to PBS or to distilled water. Small pieces about 1×2mm were cut and stained 1, 2, or 3 days in 20% lead acetate solution at room temperature (ca. 24-26°C) with some rocking or occasional agitation. Chick limbs, axolotl limbs, and mouse embryos were stained overnight in 1-10% lead(II) acetate trihydrate solution. PTA and IKI staining was carried out as reported in (Metscher, 2009a, 2009b).

For microCT imaging, samples were mounted in polypropylene pipet tips or small polyethylene tubes in 1% low-melting temperature agarose in water (Metscher, 2011). MicroCT scans were made using an Xradia MicroXCT system (Zeiss Microscopy) equipped with a tungsten X-ray source and scintillator detectors with secondary optical magnification. Source anode voltage was set to 40 or 60kV. Projection images were taken every 0.2° over a half-rotation of the sample; tomographic slices were reconstructed using the Xradia XMReconstructor software.

Images were analysed and illustrations prepared using XM3DViewer (Xradia), Fiji (Schindelin et al., 2012; https://fiji.sc/), Adobe Photoshop, and Dragonfly (https://www.theobjects.com/dragonfly).

## RESULTS

Aqueous lead(II) acetate staining was demonstrated first in mouse small intestine samples, which are known to have a distinctive pattern of cell nuclei in histological sections (Fig. 1A). Using a 20% aqueous solution (~0.53M pH ~5.5), the lead staining was found to impart X-ray contrast to cell nuclei and other tissue components, without a hematein staining step (Fig.1). Staining durations of 1, 2, and 3 days were found to be effective. The staining results with intestine samples were somewhat inconsistent, likely owing to the variable enteric environment; some high-contrast debris can be noted in Fig. 1B.

**Fig. 1:**
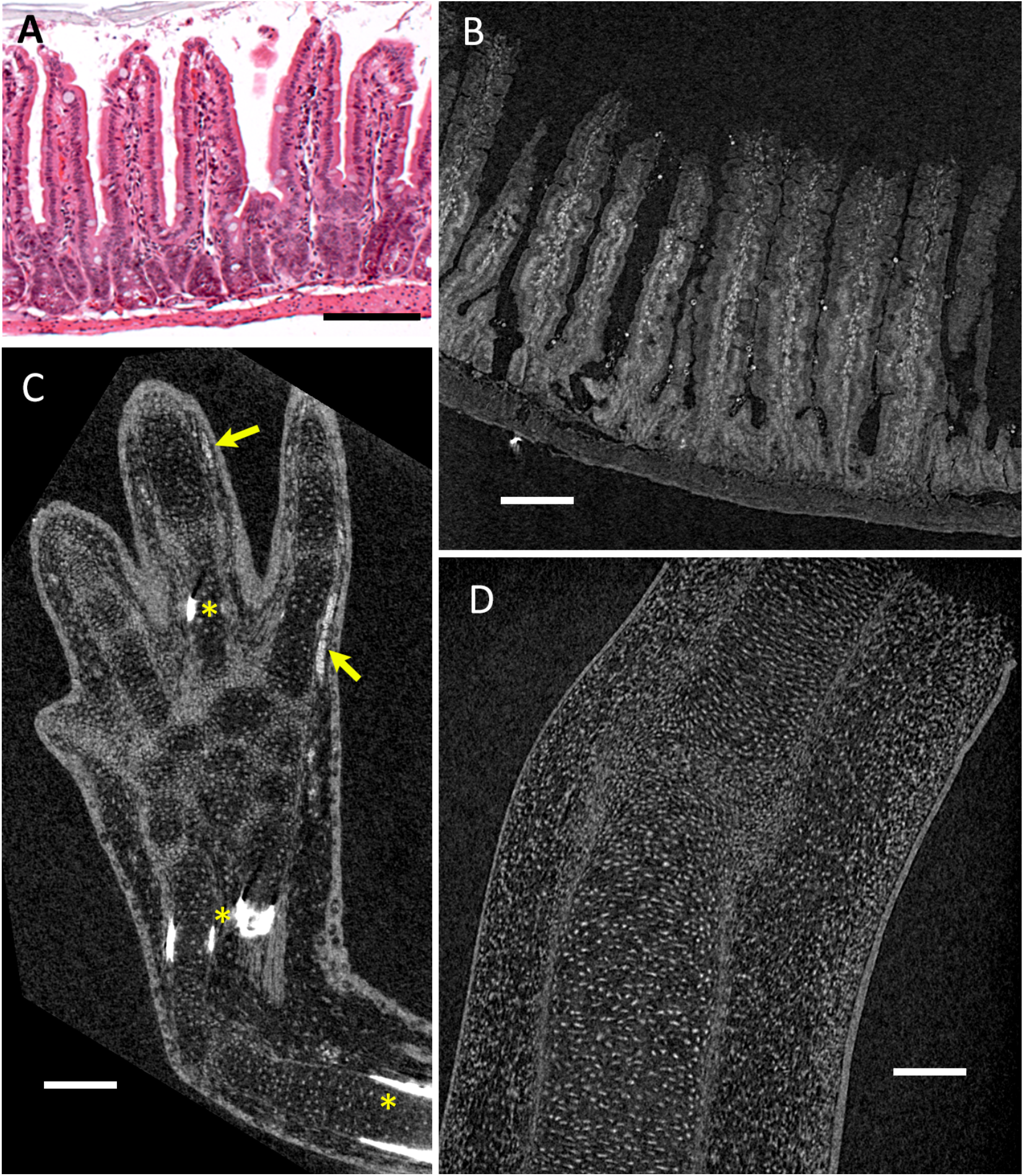
Bleikern staining highlights cell nuclei. Scale bars 100μm. A) Mouse small intestine, H&E section, reproduced from (Schweer et al., 2013). B) Adult mouse small intestine stained in 20% lead(II) acetate solution (aqueous) for 2 days at room temperature. Maximum-intensity projection of 4 consecutive virtual slices from a microCT scan; 1.0μm voxel size. C) Axolotl larva forelimb stained overnight in 2% lead(II) acetate. Single virtual section, 2.0μm voxel size. Blood cells (arrows) and early ossification of bone (stars) are indicated. D) Chick embryo (stage ~HH30) digit stained overnight in 2% lead(II) acetate. Single virtual section, 1.0μm voxel size.

Several other samples were tested to show the contrasting of cell nuclei qualitatively. Amphibians are known to have particularly large cells, and the nuclei show clearly in a larval axolotl limb (Fig. 1C). It was also observed that the early bone matrix in the limb gained much higher X-ray absorption with lead staining.

Studies of limb morphogenesis commonly require mapping of cell densities, and a chick embryo limb stained overnight allowed microCT imaging of the cell distribution (Fig. 1D). Some shrinkage was observed in lead-stained limbs. EDTA treatment (e.g. 50mM EDTA for 1 hour) removed the lead staining from tissues without apparent damage (data not shown).

A lead(II) acetate solution of 2% (~53mM) was found to stain nuclei effectively with an overnight incubation (approx. 18 hours) at room temperature (which was around 26°C during the summer these experiments were conducted). Compared with two standard microCT contrast stains, phosphotungstic acid and Lugol’s iodine (PTA and IKI; Metscher 2009a,b), the lead staining is more preferential for nuclei (Fig. 2). Bleikern stains the nuclei in embryonic blood cells (Fig. 1C), but unlike PTA and IKI, it does not stain blood more strongly than other tissues (Figs. 2B, 2C).

**Fig. 2:**
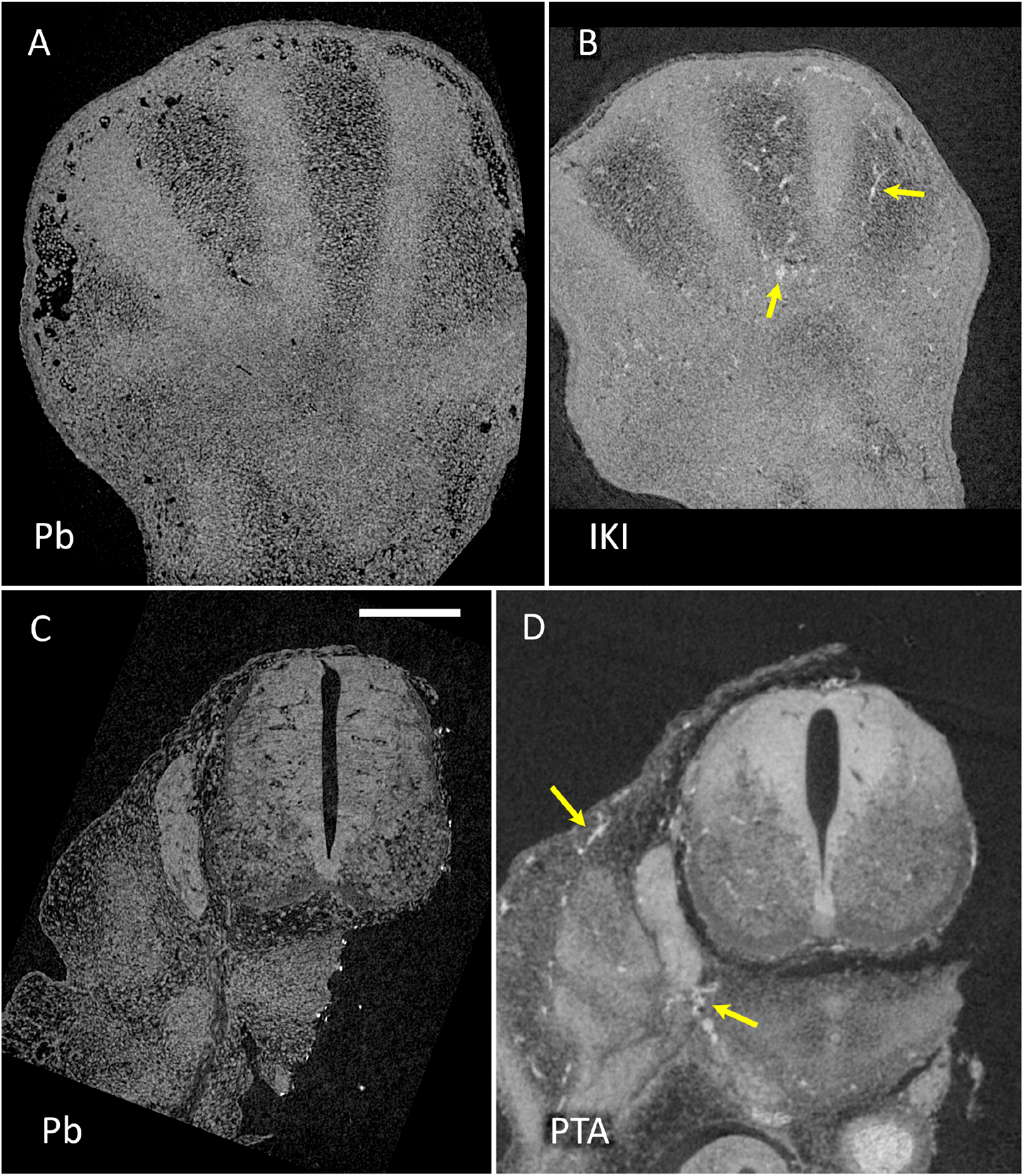
Comparison of Bleikern (overnight staining in 2% lead(II) acetate) with other stains in higher-resolution scans. Virtual sections from microCT images of mouse embryos, approximately E12.5 (Theiler stage ~21). Scale bars 200μm. A) Mouse limb, Bleikern stained. 1.0μm voxel size. B) Mouse limb IKI stained. C) Mouse spinal region, Bleikern, 1.1μm voxel size. D) Mouse spinal region, PTA stained. Arrows indicate blood.

Because microCT imaging is often employed most effectively for overview visualisation of intact samples, this staining was also tested on whole mouse embryos. A single overnight staining in 2% lead(II) acetate solution was sufficient for complete contrasting of a 12.5-day (approx. Theiler stage 21) mouse embryo, previously fixed in paraformaldehyde (Fig. 3). This method allows differentiation of tissues according to local cell density, and thus can function as a general histological stain at lower resolutions. The staining is uniform throughout the embryo and does not accumulate in the surface regions. However, X-ray dense particles were sometimes observed on the sample’s surface (e.g. Fig. 2C).

**Fig. 3:**
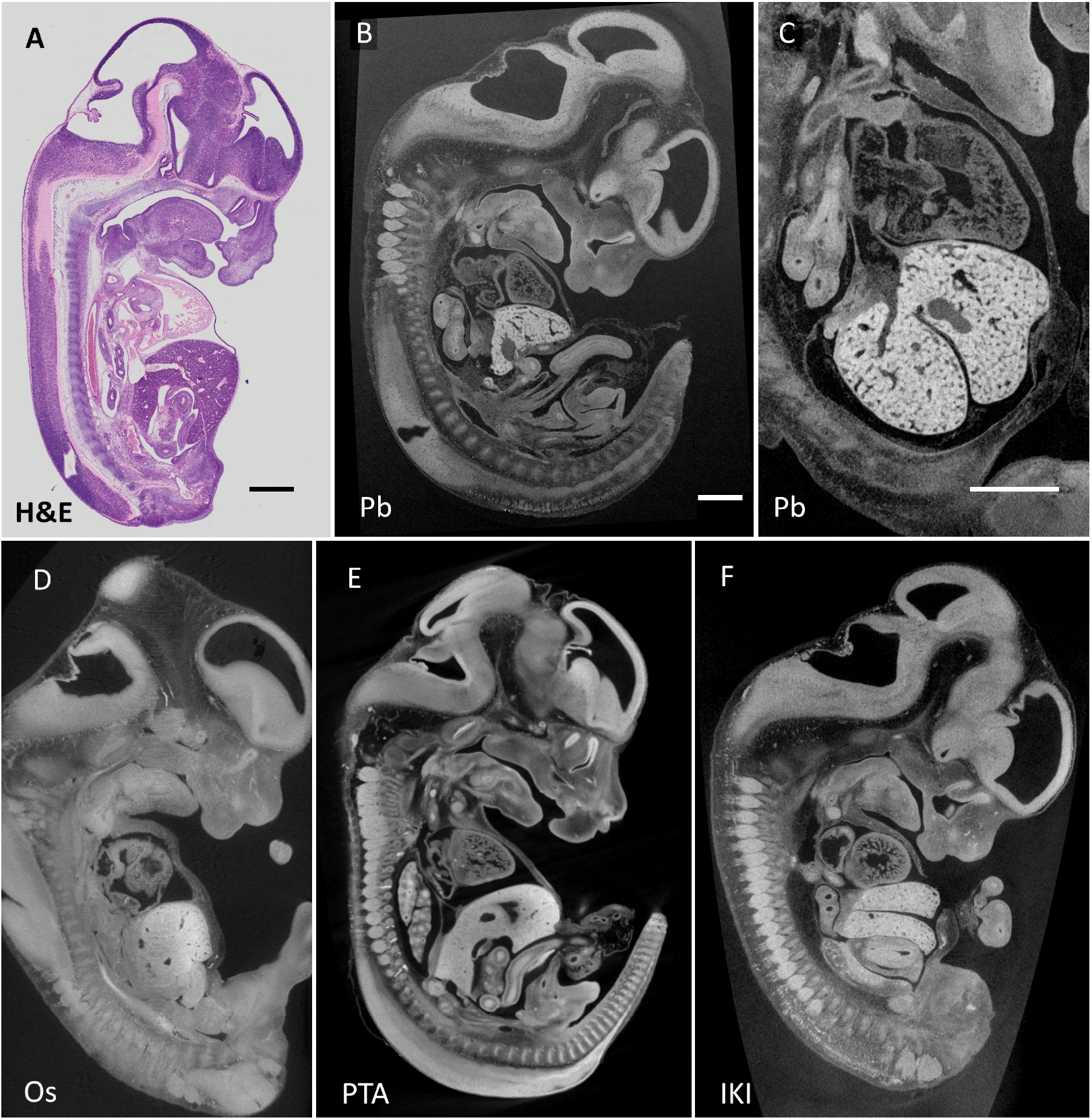
Bleikern as a general contrast stain, compared here with other stains in whole mouse embryos ~E12.5 (Theiler stage ~21). Scale bars 500μm. B-F are single virtual sections from microCT scans. A) H&E stained section from the *eHistology Atlas with Kaufman Annotations,* Plate 29a, Image a, (Graham et al., 2015; http://www.emouseatlas.org/emap/eHistology/. B) Bleikern staining, 2% overnight. Complete image data available at doi:10.5281/zenodo.4034294. C) Bleikern, same scan as in B), showing differentiation of organs according to cell density. 2.5μm voxel size. D) OsO_4_, sample prepared as for TEM (Metscher, 2020). E) PTA, favours collagen, penetrates more slowly and stains blood heavily. F) IKI, can emphasise keratin; also stains blood strongly.

## DISCUSSION

The results presented here show that an aqueous solution of lead acetate at 2% concentration (~53mM) works as a preferential nuclear stain for X-ray microtomography, but it is not highly specific to nuclei. For example the first images are not clean enough to allow segmentation of the nuclei by global thresholding of the image. These preliminary tests demonstrate some potential of the method for cell-resolution histology, though the results are limited by the useful resolution of a 14-year-old lab-based microtomography system.

The procedure demonstrated here did not involve critical-point drying, thus allowing scanning in liquid medium and leaving open the possibility of further processing of the same sample. Neither acidic re-fixation in Bouin or FAA nor acidification of the lead solution with acetic acid made any noticeable improvement to the specificity or intensity of the nuclear stain. However, it is still possible that acid treatment and hematein complexing might enhance the nuclear specificity of the staining (Kiernan, 2008b).

It was found that care was needed to avoid exposing the lead(II) to anions that cause unwanted precipitation. Thus the samples were washed in distilled water before and after staining. Subsequent dehydration to alcohol did not increase non-specific lead deposition, nor did washing in ammonium acetate solution, but neither gave any noticeable improvement to signal or background. Tap water, whose ion content can vary greatly, is likely to give unpredictable results. (Ammonium acetate solution did not cause precipitation and might offer an alternative wash buffer.)

Staining by divalent lead was easily removed with EDTA, allowing the possibility to stain and image precious samples without damage, such as museum specimens (Akkari et al., 2018, Keklikoglou et al., 2019).

In samples with early bone mineralisation, the lead staining increased the X-ray absorption of the bone matrix substantially, presumably due to Pb^2+^-Ca^2+^ ion substitution (stars in Fig. 1C). This effect could possibly be used to enhance the detection of early mineralisation of skeletal elements and teeth.

The present study was only to demonstrate a simple and fast method for contrasting cell nuclei for microCT imaging and determine its general efficacy. Quantifying the nuclear specificity and other staining effects will require experiments designed for those purposes. Sub-micron X-ray microtomography applied to standard samples such as onion roots or sea urchin embryos may provide more direct tests to determine the specificity of Bleikern staining to nuclear proteins vs. nucleic acids.

A baseline staining procedure is given here as a starting point for applying this method. The following steps must be tested and optimized for any new sample.

1. Samples fixed in formalin or paraformaldehyde.

*The effect of acidic fixatives (Bouin, FAA) has not been tested.*
2. Samples stored in fixative, buffer-diluted fixative, 70% ethanol, or methanol.
3. Take samples to distilled water.

*Soluble anions can precipitate lead(II) salts, so tap water should be avoided.*
4. Staining solution: 2% (w/v) lead(II) acetate trihydrate in distilled water.

*Like other lead compounds, lead acetate is toxic and must be handled according to standard safety procedures*. *The staining solution was still effective after two weeks at room temperature. Its maximum shelf-life was not determined*. The permanence of the stain has not been tested.
5. Incubate samples in staining solution overnight or longer with rocking or occasional agitation.

*It is possible that at least some samples can be stained for shorter times; minimum or optimal staining times and penetration rates were not assessed, and the permanence of the staining has not been tested.*
6. Change samples to distilled water and wash several times over an hour or more.
7. Samples can be moved to methanol or 70% ethanol.
8. Mount sample in agarose or alcohol for microCT imaging.

## DATA AVAILABILITY

The original tomographic image data is available on request from the author.

## ACKKNOWLEDGMENTS

The author is grateful to Nina Kraus for her assistance in carrying out this work. No conflicts of interest are associated with this work.

